# Ex Vivo Assay for Organ-Specific Cancer Cell Invasion

**DOI:** 10.64898/2026.01.08.698145

**Authors:** François Tyckaert, Paul Frieso Göddertz, Maria Reichhold, Bettina Sarg, Klaus Faserl, Pere Patón González, Felix Eichin, Andreas Villunger, Steffen Ormanns, Stefan Redl, Julia Hofmann, Theresa Hautz, Francesco Baschieri

## Abstract

**Background:** Metastasis is the leading cause of cancer-related mortality, yet experimental models often fail to recapitulate the tissue-specific microenvironments that shape metastatic dissemination. While in vivo systems provide physiological relevance, they are poorly suited for mechanistic studies. Conversely, conventional in vitro assays lack the organ-specific extracellular matrix (ECM) context that critically regulates invasive behavior. There is therefore a need for accessible models that balance biological relevance with experimental feasibility.

**Results:** In this study, we developed an ex vivo invasion platform based on mild detergent decellularization of mouse organs followed by vibratome slicing. This approach generates optically transparent lung, liver, and intestine ECM scaffolds that preserve native matrix architecture, mechanical properties, and retain biochemical hallmarks of their tissues of origin. Organ-derived matrices were integrated into standard microfluidic channels and analyzed using conventional fluorescence microscopy to enable quantitative assessment of cancer cell invasion.

Benchmarking with breast cancer cell lines of defined invasive capacity, we could demonstrate the robustness and biological relevance of the system. Non-invasive MCF7 cells failed to infiltrate any organ scaffold, whereas highly invasive MDA-MB-231 cells exhibited pronounced organ-specific invasion. These cells preferentially invaded lung and liver ECM while showing minimal invasion of intestinal scaffolds, recapitulating clinically observed metastatic tropism. Quantitative invasion rates closely matched values previously reported in vivo using intravital microscopy.

**Conclusions:** This ex vivo organ-derived ECM platform provides a scalable, cost-effective, and experimentally accessible system to study ECM-driven determinants of metastatic invasion. By preserving tissue-specific matrix cues while reducing reliance on animal models, this approach provides a powerful tool to interrogate ECM-driven determinants of metastasis and to evaluate potential therapeutic interventions.

## 1. Background

Cells within our body are embedded in a complex meshwork of proteins known as the extracellular matrix (ECM) [1], [2], [3]. Although fibrillar collagens constitute the bulk of the ECM, with collagen I alone accounting for up to 12–17% of total mouse protein mass [4], the composition, architecture, and mechanical properties of the ECM vary markedly between organs, reflecting their distinct physiological requirements [3], [5].

Beyond its structural role, the architecture and mechanical properties of the ECM guide cell movement, determine regions of tissue growth, and even dictate where cellular extrusion occurs [6], [7], [8], [9], [10]. This regulation is mediated by a finely tuned interaction between ECM components and cell adhesion molecules, such as integrins, on the cell surface [11].

The importance of cell–ECM interactions has long been recognized in cancer biology. More than a century ago, Paget’s “seed and soil” hypothesis proposed that metastatic dissemination depends on the compatibility between disseminating cancer cells and the microenvironment of target organs. Consistent with this idea, clinical and experimental evidence demonstrates that metastases arising from distinct primary tumors exhibit reproducible organ-specific dissemination patterns, a phenomenon known as organotropism [12], [13], [14]. While soluble factors and stromal cells contribute to metastatic niche formation, the ECM represents a fundamental and relatively stable component of the metastatic soil.

Metastasis is the leading cause of cancer-related mortality, and improving our understanding of this process remains a central challenge in efforts to enhance patient outcomes. Despite its importance, ECM-driven organotropism remains difficult to investigate experimentally. *In vivo* mouse models offer physiological relevance but are time-consuming, costly, and challenging to dissect mechanistically. Conversely, conventional in vitro assays are easier to interpret but fail to recapitulate the tissue-specific ECM context that critically shapes metastatic behavior, contributing to poor predictive power in preclinical studies.

There is a pressing need for improved in vitro or ex vivo models that better mimic the complexity of metastatic processes preserving the complexity of organ-specific ECMs. To address this gap, we developed an *ex vivo* invasion assay based on decellularized mouse organs. By combining mild detergent-based decellularization with vibratome slicing, we generate thin, optically transparent tissue sections that retain native ECM architecture, mechanics, and composition. These scaffolds can be integrated into standard microfluidic channels and imaged using conventional microscopy, enabling quantitative and comparative analysis of cancer cell invasion across distinct organ-derived ECMs.

## 2. Methods

### 2.1. Tissue decellularization

Immediately after collection, organs from healthy, 6-10 week old, male BL6N mice were quickly washed in PBS to eliminate excess blood. Importantly, the mice had been euthanized for unrelated procedures, and only surplus organs were used in this study. Liver lobes were separated, and intestine lumens (duodenums) were flushed multiple times with PBS. Organs were then immediately submerged in 50 ml of PBS containing 0.1% SDS and incubated rotating at room temperature (RT). The solution was changed every two days until full decellularization of the organs, which could be evaluated empirically by observing the organs through a source of light to assess their transparency. Approximately 15 days were needed to decellularize liver and lungs, while gut decellularization was achieved in 7 days.

SDS being an ionic detergent, a 3h wash with the non-ionic detergent Triton X-100 (0.05%) was used to wash away SDS, followed by 4 days washing in 50 ml H2O, with H2O changed daily. Organs were then stored in PBS until slicing. Quantification of residual SDS was performed using a colorimetric assay based on a carbocyianine dye (Stains all – Sigma-Aldrich – Cat. Nr. E9379-1G) as previously described [15]. Residual DNA was extracted from 1.5-3.5 mg of dry mass of tissue using the Tissue DNA Purification Kit (EurX – cat. Nr. E3550-01) according to manufacturer’s instructions and quantified by NanoDrop absorbance at 260 nm (260/280 ratio) in three independent replicates for each organ.

### 2.2. Tissue slicing

Decellularized organs were then included in low temperature melting agarose at 1% for liver and lung, and 3% for intestine. Organs were sliced with the help of an Alabama R&D Tissue Slicer (Alabama R&D, Munford, AL, USA) using stainless steel razor blades (Personna Medical, Stainton, VA, USA) in PBS. Native organs were sliced in ice cold PBS and slices were immediately incubated in the presence of proteases inhibitors. Slices of 500 µm were recovered in PBS and incubated at 37°C for 30 min to eliminate agarose residues, then stored at 4°C in PBS. Of note, as organs are likely to experience some minor volume changes, if the final volume of the slices is important, slicing has to be performed on already decellularized organs.

### 2.3. Stiffness measurements

A Chiaro Nanoindenter (Optics11 Life) was used to measure the mechanical properties of the tissues. Vibratome cut slices of tissues were fixed on a plastic tissue culture dish with the help of super-glue, then submerged into PBS. For the intestine, measurements were performed on the external wall of each slice. For fresh tissues, proteases inhibitors and propidium iodide were added to the PBS to prevent and monitor cellular damage. Fresh tissues were analyzed within 8 hours of collection, a time window during which minimal to no cell death was observed.

Measurements were performed with a rounded-tip probe with 27.5 µm diameter and stiffness of 0.5 N/m. Indentations of 3 µm were performed with an approach speed of 3000 nm/s. For each tissue slice, at least 36 measurements spaced of 100 µm were acquired. Curves were fitted using the Prova software from Optics11 life. The upper 1000 nm of indentation were fitted using Hertzian contact model and assuming Poisson’s ratio of 0.5 for the calculation of the effective Young’s elastic modulus.

### 2.4. Histology

Native or decellularized organs were fixed with PFA 4% for 24h and embedded in paraffin. Four micrometer thick sections of paraffin embedded tissues were stained using automated routine hematoxylin-eosin staining (Sakura Prisma, Sakura Finetek Europe, Umkirch, Germany), the connective tissue stainings Elastic van Gieson (EvG), chromotrop-anilin blue (CAB), Alcian Blue (AB), and Periodic Acid–Schiff (PAS) on an automated special staining device (Ventana BenchMark, Roche, Tucson, AZ, USA) according to the manufacturer’s instructions. Slides were scanned using a Leica Aperio GT450 DX or on a Leica epifluorescence microscope equipped with a K3C color camera and 10x objective. Acquired images were cropped and further processed with the QuPath software and FIJI.

### 2.5. Scanning Electron Microscopy

Decellularized organs were fixed in 2.5% GA and 2% PFA diluted in sodium cacodylate buffer (0.1M, pH = 7.4) for several days and postfixed in 4% Glutaraldehyde in 0.1M PB, pH 7.4 overnight. After washing with PB, samples were cryoprotected overnight in 30% sucrose, embedded in OCT and the surface was trimmed and cut using a Leica Cryocut CM3050 S. After washing in 0.1M PB, samples were postfixed in 1% OsO4 in A.d., dehydrated in a standard ethanol series and critical point dried using a BAL-TEC CPD 030 critical point dryer, mounted with carbon adhesive tape and Leit-C on aluminium stubs and sputtered with 10 nm gold using a BAL-TEC MED 020 coating system. Scanning electron microscopic images were taken with a Tescan-Clara, at the Zoology department, University of Innsbruck.

### 2.6. Proteomics

Samples were dried in a SpeedVac. Each individual sample was recovered in 40 µL of 100 mM ammonium bicarbonate (NH4HCO3), 1% deoxycholate (DOC) and 50 mM dithiothreitol and incubated at 56°C for 1 h. The samples were alkylated with 5 µl of 500 mM IAA solution for 30 min at RT in the dark. Samples were then dried again in a SpeedVac, resuspended in 50 µL of trypsin solution (10 ng/µL) in 50 mM NH4HCO3 and incubated overnight at 37°C [16]. By adding 10 µl concentrated formic acid samples were acidified (pH lower than 2) to precipitate deoxycholate. Afterwards, they were centrifuged for 10 min at 36000g and transferred to a 200 µl low binding Eppendorf tube.

For LC-MS analysis an UltiMate 3000 nano-HPLC system coupled to an Orbitrap Eclipse mass spectrometer (Thermo Scientific, Bremen, Germany) was used. The peptides were separated on a homemade frit-less fused-silica micro-capillary column (75 µm i.d. x 360 µm o.d. x 20.5 cm length) packed with 2.4 µm reversed-phase C18 material (Reprosil). Solvents for HPLC were 0.1% formic acid (solvent A) and 0.1% formic acid in 85% acetonitrile (solvent B). The gradient profile was as follows: 0-4 min, 4% B; 4-117 min, 30% B; 117-127 min, 100% B. The flow rate was 250 nL/min [17].

The Orbitrap Eclipse mass spectrometer was operating in the data dependent mode with a cycle time of one second. Survey full scan MS spectra were acquired from 375 to 1500 m/z at a resolution of 60 000, a maximum injection time (IT) of 118 ms and automatic gain control (AGC) target 400 000. The MS2 Spectrum was measured in the Orbitrap analyzer at a resolution of 15000 with an isolation window of 1.2 mass-to-charge ratio (m/z), a maximum IT of 22 ms, and AGC target of 50 000. The selected isotope patterns were fragmented by higher-energy collisional dissociation with normalized collision energy of 30%.

Data analysis was performed using Proteome Discoverer 3.1 (Thermo Scientific) with Sequest search engine. The raw files were searched against UniProt mouse-2024_UP000000589_10090 database. Precursor and fragment mass tolerance was set to 10 ppm and 0.02 Da, respectively, and up to two missed cleavages were allowed. Carbamidomethylation of cysteine was set as static modification, oxidation of Methionine, Galactosyl and Glucosylgalactosyl were set as variable modifications. Peptide identifications were filtered at 1% false discovery rate. For label free quantification the protein abundances were normalized to total protein amount.

Three biological replicates were analyzed for each decellularized organ. Only proteins detected in all three replicates were retained for further analysis. To generate organ-specific protein lists, we additionally applied a threshold of ≥0.01% of the total protein mass. Gene ontology (GO) analysis was then performed online using String database version 12.0 with the following parameters: False Discovery Rate (FDR) ≤ 1E-5; Signal ≥ 2; Strength ≥ 1; Minimum count = 10.

### 2.7. Cell culture and in vitro testing

MDA-MB-231 cells (a gift from G. Montagnac, Inst. Gustave Roussy, Villejuif, France; ATCC HTB-26). MCF7 and HEK293T cells were obtained from H. Farhan lab (source: ATCC). SW620 were purchased from ATCC. SW620 cells were lentivirally transduced with a construct encoding the plasma membrane marker CAAX–mScarlet-I3. Transduced cells were subsequently bulk-sorted to generate a stable population expressing the fluorescent reporter. All cell lines were grown in DMEM Glutamax supplemented with 10% FCS and 100 U ml−1 penicillin/streptomycin at 37°C in 5% CO2. All cell lines were used up to passage 10 and were regularly tested by PCR for the absence of mycoplasma contamination. For channel invasion experiments, Primocin (Invivogen – Cat. ant-pm-05) was also added to the medium at the final concentration of 100µg/ml. Lentiviruses were produced in HEK293T cells using the packaging plasmid psPAX2 and the VSV-G envelope expressing plasmid pMD2.G according to the Trono lab protocol. Stable cell lines expressing empty pLVTHM (a gift from Didier Trono, Addgene plasmid #12247) were obtained by lentiviral transduction, followed by sorting via fluorescence-activated cell sorting.

### 2.8. 2D cell migration

For wound healing, Ibidi 2 well culture inserts (Cat. Nr. 80209) were used. Briefly, 30.000 cells per well were seeded on uncoated plates or on collagen I coated (50 µg/ml) plates. Upon cell adhesion the inserts were removed, and cell migration across the wound was imaged (1 image every 20 min). Imaging was performed on an epifluorescence microscope (Leica Microsystems Ltd., Wetzlar, Germany) through a 10× (Fluotar N.A. 0.32 PH1) objective. The microscope was equipped with an LED5 lamp from Leica, a filter cube DFT51010, an external wheel EFW for DMi8, and an sCMOS K5 camera. Wound healing closure was analyzed using FIJI. Briefly, the area covered by cells was measured at 2, 4, 6, and 8 hours after insert removal. To account for differences in the initial wound size, data were normalized by arbitrarily setting the 8-hour time point of the “MCF7 collagen” condition to 1, and all other measurements were expressed relative to this value.

### 2.9. Gelatin digestion

For Gelatin digestion, glass coverslips were activated with the help of a plasma cleaner (Harrick Plasma PDC-002-CE). Activated coverslips were incubated for 10 min with Gelatin Oregon Green™ 488 Conjugate (Invitrogen – Cat. Nr. G13186) diluted at 1 mg/ml. Coverslips were then rinsed in PBS, and glutaraldehyde was used to crosslink gelatin, followed by quenching with Sodium Borohydride. Cells were plated and incubated at 37°C for 24h. Then, cells were fixed and stained with Phalloidin iFluor 555 (Abcam – Cat. Nr. ab176756). Images were acquired using a 60× TIRF objective (Plan-APOCHROMAT 60×/1.49 Oil, Nikon) mounted on an inverted microscope (Eclipse Ti2-E; Nikon) with a spinning disc confocal unit (X-Light V2 – 60 μm pinhole), a sCMOD camera (Photometrics Prime 95B), and Lumencor Spectra III LED light source (380 – 750 nm), controlled via the NIS-Elements software (Nikon). Cells were automatically segmented based on the phalloidin channel, and the total cell area was measured. The fluorescent gelatin channel was then thresholded to identify regions of matrix degradation, and the area of the resulting holes (corresponding to digested regions) was quantified. Results are expressed as the digested area normalized to the total cell area (percentage of degraded area per cell area).

### 2.10. Transwell invasion assays

Fluoroblok™ inserts with 8 μm membrane pores, 24 well plate format (BD Falcon – Ref. 351152) were coated with 50 μl Geltrex (Gibco – Life Technologies – Cat. Nr. A14132-02) diluted in DMEM with with 10% FCS to a final concentration of 1.2–1.8 mg/ml. Geltrex was let to polymerize for 1h at 37°C. 600 µl of DMEM supplemented with 10% FCS and 100 U ml−1 penicillin/streptomycin were added in each well, while 1E5 cells were resuspended in a volume of 300 μl of medium with 10% FCS and plated inside the coated invasion insert. Cells were then allowed to invade at 37°C for 48h. Then, cells were fixed with PFA and stained for DAPI. Imaging was performed on an epifluorescence microscope (Leica Microsystems Ltd., Wetzlar, Germany) through 63× [Numerical Aperture (N.A.) 1.40], 20× (HC N.A. 0.40 PH1), 10× (Fluotar N.A. 0.32 PH1), or 5× (PLAN 5/N.A. 0.12 PHO) objectives. The microscope was equipped with an LED5 lamp from Leica, a filter cube DFT51010, an external wheel EFW for DMi8, and an sCMOS K5 camera. A PeCon chamber was installed on the microscope to perform live imaging at 37°C with 5% CO2. The microscope was steered by the Leica dedicated LasX software with Navigator, to obtain mosaic stitched images. Using this function, the entire lower side of the Fluoroblok™ was imaged and cells having invaded were counted using FIJI. Briefly, cell nuclei were detected using the Find Maxima function on the background subtracted DAPI image, and the resulting signals were quantified with the Count Dots tool.

### 2.11. Inclusion of organs into microchannels

Ibidi sticky-Slide VI 0.4 (Cat. Nr. 80608) and polymer Coverslips for sticky-Slides (Cat. Nr. 10813) were used to host decellularized tissue slices. According to manufacturer’s instructions, channel thickness is 400 µm ± 130 µm. Organ slices were incubated in FCS for 1h prior to being included in the open sticky slide, one slice per channel, paying attention not to completely block the channel. Channels were then sealed with the help of the Ibidi Adapter for Sticky Slides (Cat. Nr. 80043) and a clamp. DMEM supplemented with 100 U ml−1 penicillin/streptomycin and Primocin (Invivogen – Cat. Nr. ant-pm-05) at 100 μg/ml was added to the channels, and organs were let equilibrate in medium for at least 3h at 37°C in 5% CO_2_. 1×10⁵ cells were resuspended in 30 µl DMEM with antibiotics and injected in the channels. Slides with organs and cells were let upside down under agitation at RT for 30 min, then non adherent cells were washed away and 100 µl of DMEM medium supplemented with antibiotics and FCS were added in each channel. Slides were then incubated at 37°C in 5% CO_2_ for the whole duration of the experiment, replacing medium every other day.

### 2.12. Immunostaining and imaging

PFA was added in the channels for 10 min, followed by washes with PBS. Collagen IV antibody was purchased from Abcam (Cat. Nr. Ab6586) and used according to the manufacturer’s instructions. Cleaved collagen antibody (ImmunoGlobe – Cat. Nr. 0217-025) was diluted 1:100 in PBS Tx 0.3% BSA 1%, added to the channels, and incubated ON at 4°C. Secondary antibodies, as well as phalloidin (Phalloidin-A647 – Invitrogen – Cat. Nr. A22287) and DAPI were then used prior to performing imaging.

Imaging was performed on a Leica Sp8 confocal microscope equipped with a fast resonant scanner (8000 Hz), an incubation chamber, two hybrid and three PMT detectors, using a white laser to excite all the fluorophores at the required wavelength. Imaging was performed using a 63× glycerol immersion objective (1.4 NA) or a 20x glycerol immersion objective (0.75 NA). Interference reflection microscopy (IRM) images were acquired using a 633 nm laser, enabling direct visualization of fibrillar ECM without additional staining. 3D reconstructions of images were prepared using Imaris Version 11.0.

### 2.13. Quantification of organ invasion

To quantify cancer cell invasion across organs, cells were allowed to invade for 7 days, after which samples were fixed and analyzed by microscopy. A cross-sectional image of each tissue slice was captured to visualize invasive foci. For each region exhibiting cellular signal within the tissue, high-resolution volumetric images were collected using a z-step of 1 µm over a total depth of 140 ± 40 µm. This step ensured that detected cells were embedded within the matrix rather than adhering to surface irregularities (e.g., invaginations). Only cells fully enclosed within the tissue were considered invasive. Volumetric image stacks were analyzed in FIJI. Invasion depth was quantified by measuring the perpendicular distance from the center of each invading cell to the nearest tissue surface. The mean of these measurements was calculated to define penetration depth within each invaded region across all organ scaffolds. Importantly, the quantification reflects the depth of invasion within invaded regions only and does not account for the total number of invasion foci per organ. This approach was chosen to minimize biases arising from differences in the size and shape of the tissue pieces used in the assay.

### 2.14. Statistical analysis

All images were analyzed in FIJI. Analysis was performed manually by two independent users, and results were independently verified to ensure concordance. All images were background subtracted prior to performing any analysis. Graphics were prepared with Prism V10.2.3 and statistical analyses were performed using Prism V10.2.3. Box-plot are built as follows: The line within the box corresponds to the mean value of at least three independent experiments (n indicated in each figure legend), boxes indicate interquartile ranges (Q1 to Q3), whiskers extend to min and max values, and, eventually, outliers are represented as points outside the box-whiskers. Empty circles on top of the graphs represent the mean of each individual experiment unless otherwise stated.

## 3. Results

### 3.1. Decellularization of mouse organs

Several decellularization strategies exist, but detergent-based methods are generally considered the most effective strategy to remove cells from tissues [18]. Recently, it was shown that using low concentration of SDS allowed to better preserve tissue structure [19]. Given the importance of mechanical and structural properties of the ECM in cancer progression [20], we hence employed SDS as a decellularizing agent.

Organs from overnumbered adult, wild type mice were hence decellularized using SDS at low concentration. Efficient decellularization was visually confirmed by loss of pigmentation and transparency, making it possible to see blood vessels with the naked eye (Fig. 1A). To ensure full decellularization, DAPI stainings were performed on sections of organs, and fluorescence quantification confirmed the absence of nuclei from decellularized organ slices (Suppl. Fig. 1A-B). Residual DNA is a common indicator of decellularization efficiency, with values below 50 ng DNA per mg of tissue dry mass generally considered a benchmark for successful cell removal [21]. Measurements of residual DNA confirmed that decellularization was effective in all three organs, with values at or near this threshold (Suppl. Fig. 1C). Additionally, hematoxylin and eosin stainings were performed on paraffin embedded slides to search for remaining cells. Visual inspection of the slices confirmed that full decellularization was achieved in all tissues (Fig. 1B). At the macroscopic level, decellularized organs retained their overall shape, suggesting that the decellularization procedure did not markedly affect tissue topography or gross mechanical properties. To further assess structural preservation, we imaged decellularized organs by scanning electron microscopy (SEM), which revealed intact microanatomical features characteristic of each tissue type (Fig. 1C). For example, we could detect the Glisson’s triad in the liver, bronchial structures in the lung, and intact blood vessels within the gut submucosa (Fig. 1D). Consistent with these observations, histological stainings for elastin and collagen performed on native and decellularized tissues confirmed that the protocol did not substantially alter the underlying tissue architecture (Suppl. Fig. 2A-B). As SDS can deplete glycosaminoglycans (GAGs) and other glycoproteins from the ECM [21], we evaluated the preservation of these components after decellularization using Alcian Blue staining, which specifically labels acidic polysaccharides such as GAGs. The staining revealed that, although some depletion occurred, a substantial fraction of glycoproteins was retained within the matrix (Suppl. Fig. 2C).

**Figure 1.**
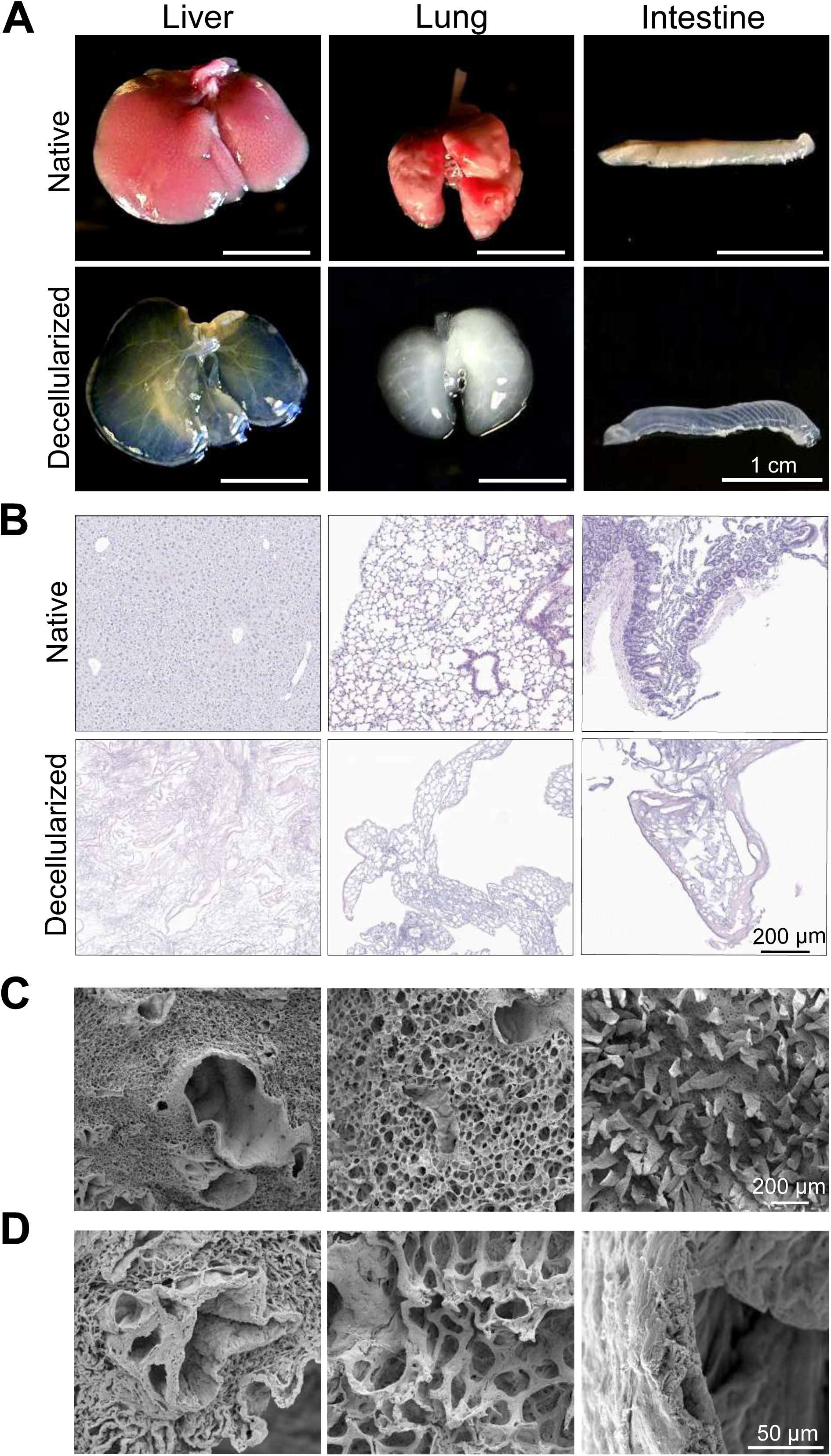
– Preparation and characterization of decellularized organs. **(A)** Representative images of native and decellularized organs. **(B)** Hematoxylin and eosin staining of native and decellularized organs **(C)** Representative SEM images of decellularized organs. **(D)** Close-up of organ-specific structural features (from left to right: Glisson’s triad for the liver, bronchi in the lung, and blood vessels included in the intestine submucosa).

Finally, as SDS is toxic to cells, optimal removal of SDS was assessed with a colorimetric assay previously described [15] (Suppl. Fig. 2D). Of note, residual SDS below 0.005% is non-toxic to human cells [22], and the washing protocol used achieved this level in all organs after just two washes.

### 3.2. Mechanical tissue properties are maintained after decellularization

We next assessed whether decellularization altered the mechanical properties of the tissues. Elastic modulus measurements were performed using a Chiaro nanoindenter (Optics11 life) on both fresh and decellularized tissue slices of lungs, liver, and gut. In this system, an optical fiber detects the phase shift caused by cantilever deflection during indentation, allowing precise measurement of tissue mechanical properties.

Slices of 500 µm of thickness were immobilized on the bottom of a petri dish and indentation measurements were performed with a round tip as described in the materials and methods. Importantly, all the tissues were analyzed using the same settings, allowing for direct comparison of the results. Native tissues were imaged in the presence of proteases inhibitors to reduce tissue damage, and propidium iodide was added to the imaging medium to visually confirm the absence of damaged cells from the measured areas. Measurements were performed over areas of 600 µm x 600 µm. No significant differences were found between native and decellularized tissue for any of the organs tested (Fig. 2).

**Figure 2.**
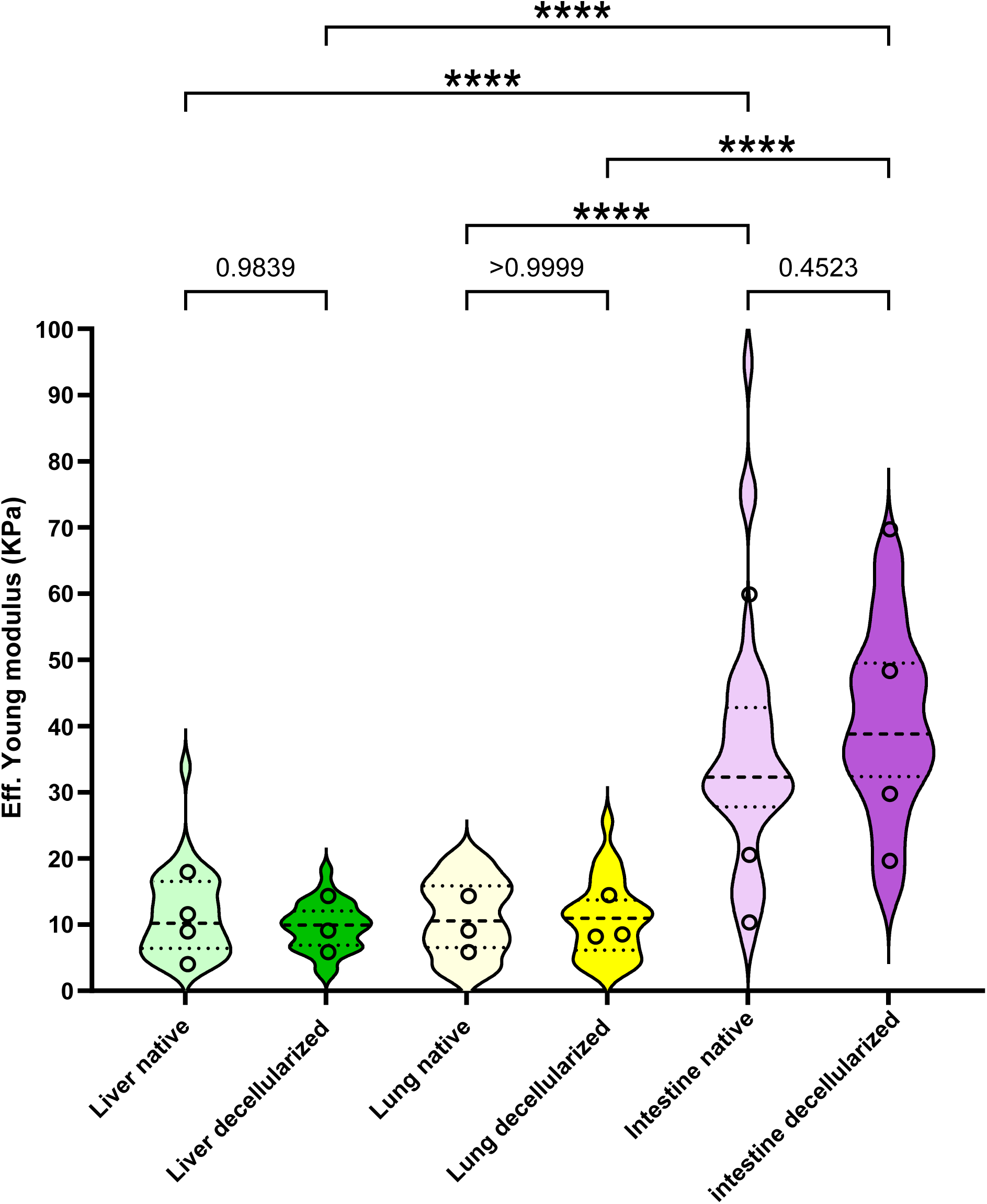
– Mechanical characterization of sliced organs. Measures of the Effective Young Modulus of slices of native or decellularized organs. 36 measures per sample were performed and violin plots were built using individual measures. Empty circles represent the average Effective Young Modulus of every organ measured. At least 3 slices, each corresponding to a different organ, were measured in each category. Results were analyzed by ANOVA Tukey’s multiple comparisons test (p values are indicated for non-significant differences; **** p < 0.0001).

These data confirm that our detergent-based decellularization procedure did not significantly alter tissue stiffness, making the decellularized scaffolds suitable for the study of mechanosensitive processes such as cell invasion.

### 3.3. Composition of decellularized organs

To evaluate the molecular integrity of the ECM, we performed label-free, quantitative proteomics on decellularized tissues (Suppl. Table 1). A core set of ECM proteins was consistently detected across all samples, along with distinct tissue-specific components. Proteins known to be enriched in each organ, such as Collagen IV in liver, Thrombospondin and Elastin in lung, and Mucins in the gut [3], [23], were reliably identified by this approach (Fig. 3A, Suppl. Table 1). Notably, these findings were corroborated by immunostaining, which confirmed the presence of elastin in lung (Supplementary Fig. 2A), Collagen IV in liver and intestine (Supplementary Fig. 3A), and partial retention of mucins in the intestine (Supplementary Fig. 2C and 3B). Gene Ontology (GO) analysis placed ECM and ECM-associated functions among the top three molecular categories in all tissues examined (Fig. 3B, Suppl. Table 1). In addition, the UNIPROT keywords “basement membrane” and “collagen” were enriched with high confidence across all decellularized scaffolds (Suppl. Table 1).

**Figure 3.**
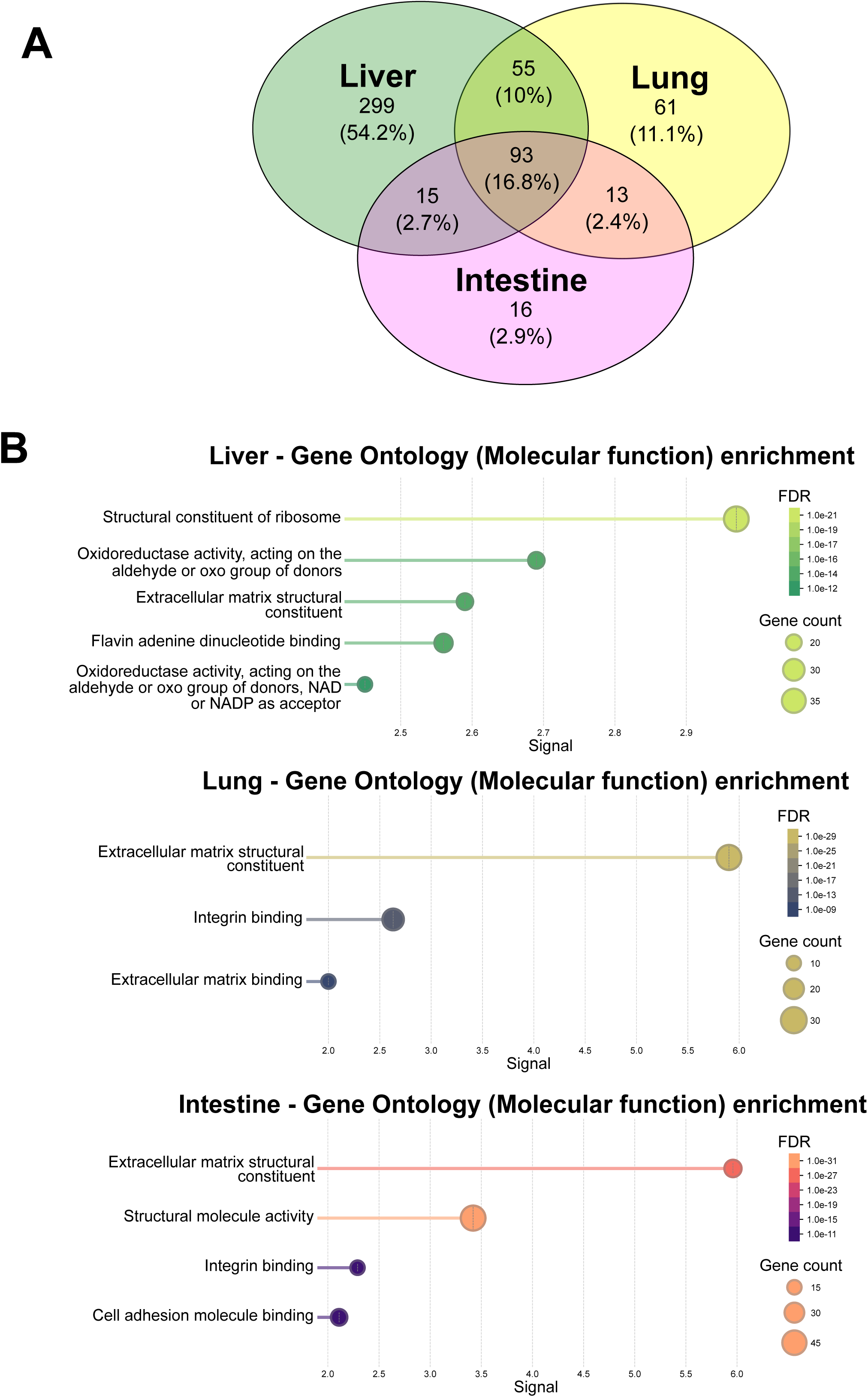
– Proteomics analysis on decellularized organs. **(A)** Decellularized organs were subjected to mass spectrometry. The number of organ-specific proteins, as well as the number of proteins shared between different organs is displayed. **(B)** Gene Ontology enrichment of the proteins identified in each decellularized organ with the (top 5) significant molecular functions listed.

Although ECM and ECM-associated proteins were strongly enriched in all decellularized organs, proteomic analysis of liver also detected some intracellular proteins despite evidence of successful decellularization, as indicated by absence of nuclei (Fig. 1B, Suppl. Fig. 1A-B) and residual DNA below 50 ng per mg of tissue dry mass (Suppl. Fig. 1C). Electron microscopy imaging of decellularized organs revealed the presence of small vesicles adhering to the liver ECM, with sizes consistent with those of various types of extracellular vesicles [24] (Suppl. Fig. 3C-D).

Together, these findings indicate that, although some loss of biochemical information occurs, our protocol largely preserves organ-specific differences in ECM composition.

### 3.4. In Vitro invasion assays confirm intrinsic cell line differences

To benchmark our ex vivo model, we first tested the invasive potential of MDA-MB-231 and MCF7 breast cancer cells in conventional 2D and 3D in vitro assays. In 2D wound healing, both cell lines displayed the ability to move, but MDA-MB-231 were significantly faster than MCF7 both on uncoated substrates and substrates coated with collagen I (Fig. 4A-B). Invasion through artificial ECM was assessed using fluorescence blocking transwells. In agreement with the literature, MDA-MB-231 were strongly invasive, contrary to MCF7 (Fig. 4C-D). The ability to degrade the matrix is necessary to invade through dense matrices. We therefore measured the degradation capacity of the two cell lines by plating them on coverslips coated with fluorescent gelatin and allowing them to degrade the substrate overnight. Again, in agreement with the literature, MCF7 did not display any degradative capacity, while MDA-MB-231 degraded the fluorescent gelatin in regions resembling focal adhesions (Fig. 4E-F).

**Figure 4.**
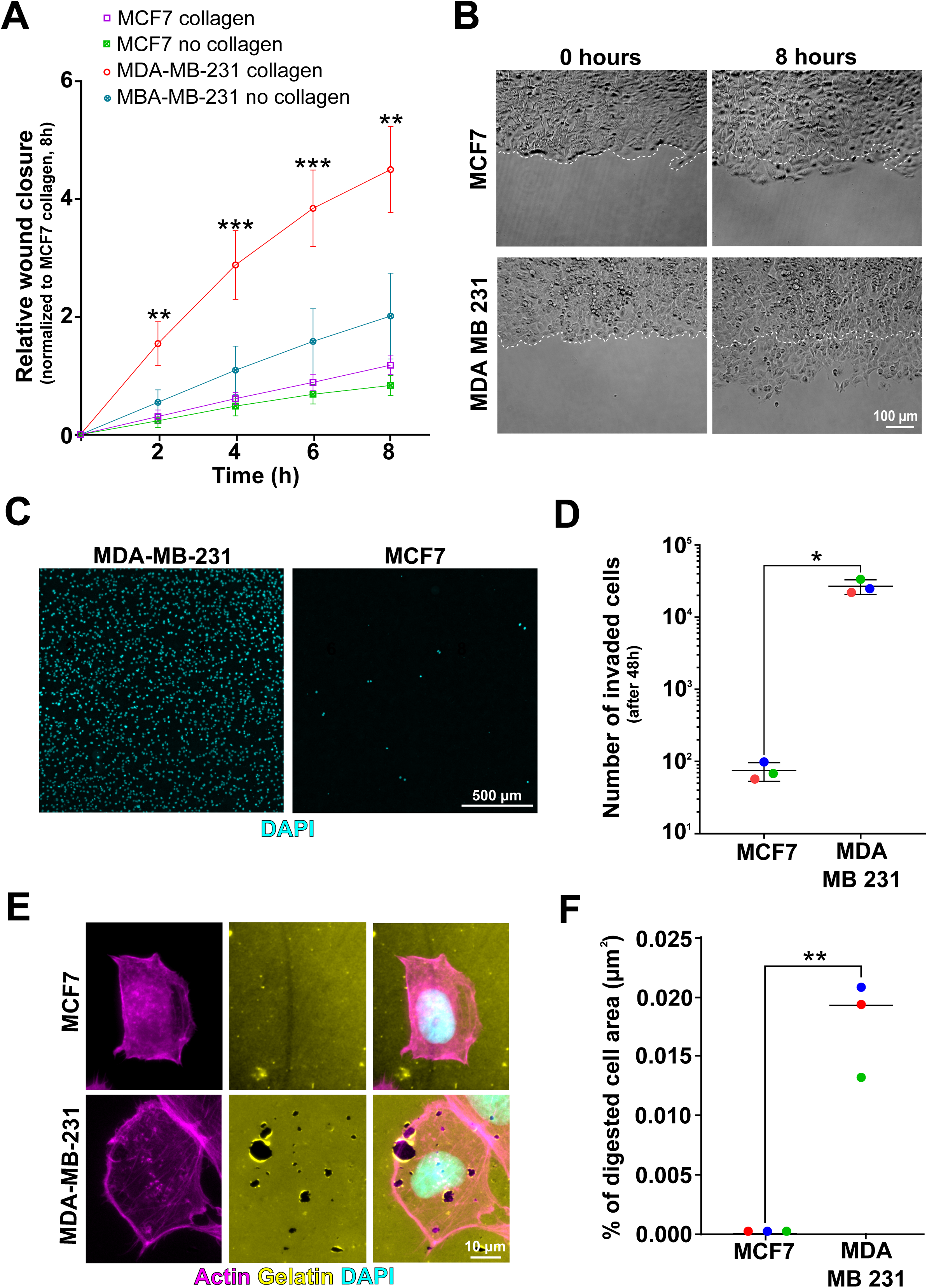
– In vitro characterization of breast cancer cell lines. **(A)** Cells were seeded to confluence in ibidi wound healing inserts. Inserts were removed and cells were imaged every 2h for a total of 8h. 5-6 wounds per cell type from 2 independent experiments were quantified. Data were normalized to MCF7 on collagen at 8h in Experiment 1. Results were analyzed by two-way ANOVA with multiple comparisons test (** p < 0.01; *** p < 0.001; statistics refer to the comparison MDA-MB-231 collagen vs MCF7 collagen). **(B)** Representative images of wounds of cells on collagen-coated substrates at the beginning (0 hours) and at the end (8 hours) of the videos. Dashed lines represent wound edges at 0h **(C)** Representative images of cells after 48 h of invasion, stained with DAPI. **(D)** Cells were allowed to invade through GelTrex for 48h. Then, invading cells were labeled with DAPI, imaged and counted. Three independent experiments, each containing 2 technical replicates per condition were performed and analyzed by Student’s t-test (* p < 0.05). **(E)** Representative images of cells seeded on fluorescent gelatin-coated coverslips, allowed to adhere for 24h, then fixed and stained for actin (phalloidin) and nuclei (DAPI). **(F)** The digested area underneath the cells (regions devoid of gelatin) was measured as compared to the total cell area (segmented based on the actin cytoskeleton) to obtain percentages of digested areas. 167 and 80 cells respectively for MCF7 and MDA-MB-231 from 3 independent experiments were analyzed and statistical differences were assessed by unpaired t-test (** p < 0.01).

These results confirm the differential invasive behavior of the two cell lines and provide a reference for their behavior in the organ-derived scaffolds.

### 3.5. Breast cancer cell invasion is organ-specific in decellularized scaffolds

For ex vivo invasion assays, organ slices were inserted into ibidi channels. Thanks to the precision of the cutting process [25], we were able to place the slices into the channels without the need for glue or excessive compression. GFP-expressing cancer cells were added and allowed to invade for 7 days, then invasion was quantified (Suppl. Fig. 4A-B). Although some autofluorescence is always present when imaging tissues, the decellularization protocol renders tissues virtually transparent and makes imaging possible using a standard fluorescence microscope without requiring fixation or staining (Suppl. Movie 1). Visual inspection of all organs, along with live imaging of lung slices, confirmed that both cell lines adhered to the scaffold surfaces from the start of the assay, ensuring that differences in invasion were not due to attachment efficiency (Suppl. Movie 1).

After 7 days of invasion, cells and tissues were fixed and subjected to immunostaining. Consistent with epifluorescence imaging, confocal microscopy with volumetric acquisition was performed to ensure that cells were truly embedded within the ECM rather than merely residing on the tissue surface. These volumetric data confirmed that MCF7 cells failed to infiltrate any of the three tissues tested. In contrast, MDA-MB-231 cells invaded the lungs and liver, whereas their penetration into the intestine was negligible (Fig. 5A-C; Suppl. Movies 2-4). To confirm that the intestine was not inherently impermeable to cellular infiltration, we tested the invasive potential of the colorectal cancer cell line SW620. These cells were able to invade all three organs tested, including the intestine (Suppl. Fig. 5A-C).

**Figure 5.**
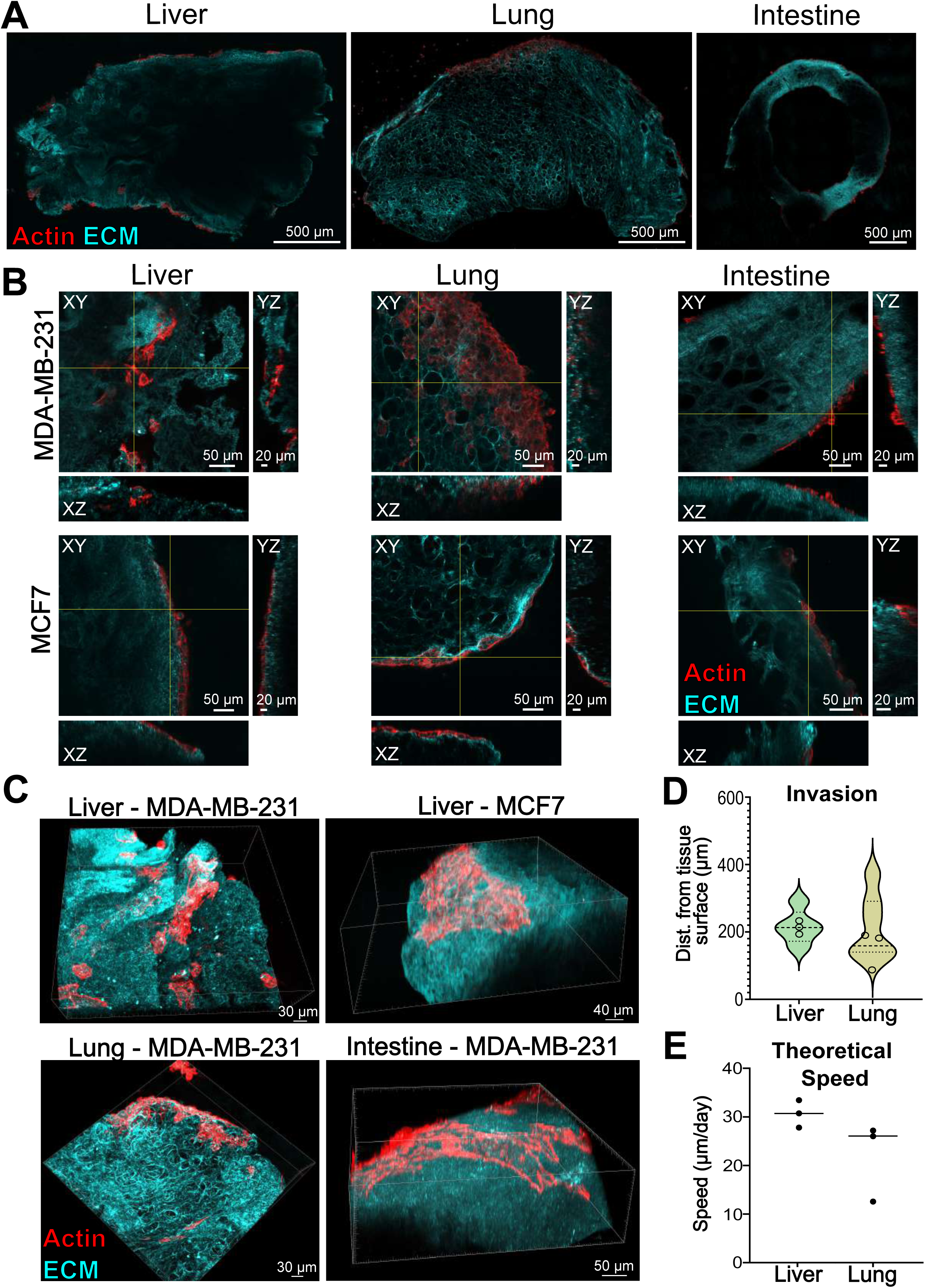
– Ex vivo invasion of breast cancer cells through decellularized organs. **(A)** Representative mid-section of organ slices after 7 days of incubation with cancer cells (MCF7 for liver, MDA-MB-231 for lung and intestine), stained with phalloidin (red). Interference reflection microscopy was used to visualize fibrillar ECM (cyan). **(B)** Orthogonal views of representative volumetric images of MDA-MB-231 and MCF7 incubated with the indicated organs for 7 days and stained for phalloidin (red). Interference reflection microscopy was used to visualize fibrillar ECM (cyan). **(C)** 3D rendering of cells in different organs. **(D)** The distance of the invasive front of MDA-MB-231 cells was quantified in the indicated organs across three independent experiments. Violin plots show invasion distances measured at multiple points along individual invasive foci for each organ. Empty circles indicate the mean invasion distance per organ. **(E)** A theoretical invasion speed of MDA-MB-231 in the indicated organs was calculated by dividing the invaded distance as in (D) by the number of invasion days.

Both single cell invasion and collective invasion could be observed in invaded organs (Fig. 5B-C). Additionally, localized collagen degradation, indicative of metalloprotease-dependent mesenchymal invasion, could be readily detected following immunostaining with an antibody specifically detecting matrix metalloprotease (MMP)-cleaved collagen (Suppl. Fig. 5D).

Quantitative analysis of invasion depth of MDA-MB-231 cells showed only a slight difference between liver and lung scaffolds, which did not reach statistical significance (Fig. 5D). Penetration distances could be converted into an average effective invasion speed, yielding values of approximately 27–30 µm/day in liver and 12–27 µm/day in lung (Fig. 5E).

Overall, these findings demonstrate that decellularized organ scaffolds quantitatively recapitulate clinically observed metastatic tropism, with breast cancer cells preferentially invading lung and liver over intestine tissue.

## 4. Discussion

In summary, we have established a versatile ex vivo invasion platform using decellularized organ scaffolds that enables quantitative, organ-specific analysis of cancer cell invasion in physiologically relevant ECM. Designed to bridge reductionist in vitro assays and complex *in vivo* models, it supports side-by-side comparisons of multiple organ ECMs prepared with a unified protocol.

Despite the recognized role of ECM in metastasis [14], such comparisons remain limited. Our data suggest that major tissue-specific mechanical and biochemical features are preserved after decellularization. However, nanoindentation reflects bulk stiffness and may miss microscale heterogeneity or subtle, decellularization-induced changes, and ECM preservation was inferred rather than directly validated against native proteomes [3].

Proteomics identified intracellular components in liver scaffolds despite evidence of effective decellularization. Electron microscopy revealed matrix-adherent vesicles resembling prior observations on decellularized ECM [26]. As the liver both receives and produces extracellular vesicles [27], [28], their retention within the matrix is plausible and may contribute to the observed proteomic signatures. However, the identity and functional relevance of these vesicles remain to be established.

Functionally, the system discriminated between invasive and non-invasive cells and revealed organ-dependent invasion behaviors not captured by conventional assays. MDA-MB-231 cells invaded lung and liver scaffolds but showed negligible penetration into intestine ECM, consistent with reported metastatic organotropism in triple-negative breast cancer [24], with invasion rates comparable to in vivo measurements [25].

Limited invasion into intestine ECM may reflect its enrichment in fibrillar collagens (notably collagen I and III) and relatively low laminin content, yielding a dense matrix with higher stiffness than lung and liver. On one hand, these ECM properties may impose physical constraints; on the other hand, differences in integrin and MMPs repertoires across cell types may further modulate matrix engagement, restricting breast cancer cell invasion while being permissive for colorectal-derived SW620 cells.

Overall, this system supports a role for native ECM in shaping invasion and is compatible with imaging and 3D reconstruction while preserving tissue architecture. The workflow is scalable and adaptable to diverse tissue sources, including mouse models of fibrosis and human biopsies. While it does not capture all aspects of the metastatic niche (e.g., immune surveillance, perfusion), it deliberately isolates ECM-specific contributions and complements 3D spheroids, organ-on-chip platforms, and in vivo assays for studying the complexity of metastasis.

## 5. Conclusions

Metastatic dissemination is shaped by organ-specific ECM cues that remain difficult to interrogate with existing experimental models. The ex vivo organ-derived ECM platform presented here establishes a standardized and accessible framework to study these cues across different tissues under controlled conditions. By enabling systematic, cross-organ analyses within preserved native matrices, this approach opens new opportunities to dissect ECM-driven mechanisms of invasion, compare tissue-specific microenvironments, and evaluate ECM-targeted interventions. More broadly, this methodology provides a foundation for developing more predictive and physiologically grounded models of metastatic behavior.

## 7. Declarations

### Ethics approval and consent to participate

Not applicable

### Consent for publication

Not applicable

### Availability of data and materials

All data generated or analyzed during this study are included in this article (and its supplementary information files).

### Authors’ contributions

**FT**: conceptualization, data curation, investigation, methodology, validation, visualization, review and editing. **PFG**: investigation, methodology, visualization, review and editing. **MR**: methodology, investigation. **BS**: methodology, investigation. **KF:** methodology, investigation. **PPG**: data curation, formal analysis. **FE**: resources. **AV**: resources. **SO**: investigation, resources. **SR**: investigation, resources. **JH**: investigation, methodology, resources. **TH**: methodology, resources. **FB**: conceptualization, data curation, formal analysis, funding acquisition, investigation, methodology, project administration, supervision, visualization, writing original draft

## 6. List of abbreviations

ECM: Extracellular Matrix
PBS: Phosphate-buffered saline
SDS: Sodium dodecyl sulfate
DMEM: Dulbecco’s Modified Eagle Medium
SEM: Scanning electron microscopy
GO: Gene Ontology
GFP: Green fluorescent protein
CAB: Chromotrop-Anilin Blue
PAS: Periodic Acid Schiff
IRM: Interference Reflection Microscopy
MMPs: Matrix Metalloproteinases

## Acknowledgements

We thank the Biooptics facility and the Proteomics facility of Medical University Innsbruck for technical support throughout the research project. We thank Dr. Anna Seybold from the electron microscopy facility of the University Innsbruck for technical support.

## Supplementary Material

**Supplementary Movie 1**

MCF7 (left) or MDA-MB-231 (right) cells were allowed to adhere to lung slices for 24h, then imaged on an epifluorescence inverted microscope at the frequency of 1 image every 10 min for 15h. Cells are GFP-positive, and decellularized organs are visualized by phase contrast. Scale bars: 50 µm.

**Supplementary Movie 2**

3D rendering of MDA-MB-231 cells adhering on the external surface of the intestine without invading. Red: cells. Cyan: Fibrillar ECM imaged by interference reflection microscopy.

**Supplementary Movie 3**

3D rendering of MDA-MB-231 cells invading into the liver. Red: cells. Cyan: Fibrillar ECM imaged by interference reflection microscopy.

**Supplementary Movie 4**

3D rendering of MDA-MB-231 cells invading into the lungs. Red: cells. Cyan: Fibrillar ECM imaged by interference reflection microscopy.

**Supplementary Table 1**

Proteomics analysis and permalinks to Gene Ontology Enrichment Analyses.

**Suppl. Fig. 1 – Characterization of the decellularization protocol (A)** Representative images of native or decellularized organs stained with DAPI. **(B)** DAPI fluorescence of images as in (A) was quantified over a surface of at least 4 mm^2^ for each organ. Results were analyzed by non-parametric LSD Fisher’s test (* p < 0.05; **** p < 0.0001). **(C)** Residual DNA in decellularized organs was measured in 3 biological replicates. No significant deviation from the 50 ng/mg reference value for successful decellularization (blue dashed line) was detected using One-sample t-test.

**Suppl. Fig. 2 – Histology of native and decellularized organs (A)** Connective tissue in the different organs was visualized by Elastic van Gieson staining. **(B)** Collagen was visualized in the different organs by Chromotrop-Anilin Blue (CAB) staining. **(C)** Glycosaminoglycans (GAGs), acidic mucins, and proteoglycans were visualized in the different organs by Alcian Blue staining. **(D)** Residual SDS in the solution where decellularized organs were washed and stored was quantified in three to five independent experiments. The blue dashed line corresponds to the maximum tolerated levels of residual SDS for recellularization purposes.

**Suppl. Fig. 3 – Validation of proteomics hits (A)** Decellularized slices of each organ were stained for Collagen IV and imaged by confocal microscopy. Interference Reflection Microscopy (IRM) was used to visualize the ECM fibers. **(B)** Native and decellularized organs were subjected to immunohistochemistry to assess the levels of Mucins (Periodic Acid Schiff – PAS) in the different organs before and after decellularization. **(C)** Scanning Electron Microscopy image of extracellular vesicles (magenta) adhering to the ECM of the decellularized liver. **(D)** Size distribution of the extracellular vesicles as in (C).

**Suppl. Fig. 4 – Schematic of the ex-vivo system’s setup (A)** Schematic of the protocol to place decellularized organ slices in imaging microchannels divided into the two phases of preparation and invasion experiment. **(B)** Cancer cell invasion was quantified as illustrated in the schematic: First, a cross-sectional image of each tissue slice was captured to visualize invasive foci. For each invaded region, a higher-resolution volumetric image was then acquired. Within these images, the distance between invading cancer cells (red) and the original organ surface (dashed black line) was measured by drawing perpendicular lines (yellow). The mean of these measurements was used to define the penetration depth of cancer cells within each invaded region across all organ scaffolds.

**Suppl. Fig. 5 – Characterization of ex vivo invasion in the different organs (A)** Orthogonal views of representative volumetric images of SW620 stably expressing the plasma membrane marker CAAX-mScarlet incubated with the indicated organs for 7 days. Interference reflection microscopy was used to visualize fibrillar ECM (cyan). (**B**) Violin plots show invasion distances measured at multiple points along individual invasive foci for each organ (n=1 for liver and lung; n=2 for intestine). **(C)** Theoretical invasion speed of SW620 in the indicated organs was calculated by dividing the average invaded distance across entire organs by the number of invasion days. **(D)** MDA-MB-231 were allowed to invade a decellularized intestine for 7 days. Then, invaded organs were fixed and immunostained for phalloidin (red) and cleaved collagen (yellow). Interference reflection microscopy was used to visualize fibrillar ECM (cyan).

